# Characterization of GRK5 as a novel regulator of rhabdomyosarcoma tumor cell growth and self-renewal

**DOI:** 10.1101/839340

**Authors:** Thao Pham, Kristin Robinson, Terra Vleeshouwer-Neumann, James E. Annis, Eleanor Y. Chen

## Abstract

Rhabdomyosarcoma (RMS) is the most common soft-tissue pediatric sarcoma. Treatment options remain limited, presenting an urgent need for novel therapeutic targets. Using a high-throughput siRNA screen against the human kinome, we identified GRK5, a G-protein receptor kinase, as a novel regulator of RMS tumor cell growth and self-renewal. Through functional assays *in vitro* and *in* vivo, we show that GRK5 regulates cell cycling in a kinase-independent manner to promote RMS tumor cell growth. GRK5 interacts with NFAT to facilitate autoregulation of *NFAT1* expression in a kinase independent manner, and loss of NFAT1 phenocopies GRK5 loss-of-function effects on cell cycle arrest. Self-renewal of RMS, required for recapitulation of tumor heterogeneity, is significantly reduced with loss of GRK5 due to increased cell death. Treatment of human RMS xenografts in mice with CCG-215022, a GRK5-selective inhibitor, reduces tumor growth of RMS. GRK5 represents a novel therapeutic target for the treatment of RMS.

**Statement of Significance:** GRK5 promotes growth and self-renewal of RMS, thereby representing a novel therapeutic target for improving survival outcomes of RMS patients. GRK5 regulates RMS tumor cell growth in a kinase-independent manner through direct interaction with NFAT1. This finding promises novel drug design, targeting non-kinase domains of GRK5.

## Introduction

Rhabdomyosarcoma (RMS) is the most common pediatric soft-tissue cancer. There are two major subtypes of RMS, each with distinct histologic features and genetic alterations. Embryonal rhabdomyosarcoma (ERMS) typically harbors mutations in the *RAS* pathway^1^. Alveolar rhabdomyosarcoma (ARMS) is characterized by the presence of the PAX3- or PAX7-FOXO1 fusion^2^. While the prognosis is good for patients with localized disease, the survival rate for patients with relapsed RMS is only 10-30%^3^, highlighting an urgent need for more effective treatment options for disease relapse. Tumor propagating cells (TPCs) are thought to be responsible for metastasis and relapse of some cancer types, such as breast and lung cancer^4–7^, and possess stem cell-like characteristics that allow for the recapitulation of tumor heterogeneity in its entirety^7^. A potential TPC population with self-renewal capacity has been identified in a conserved transgenic zebrafish model of ERMS^8^. In human ERMS, CD133-positive cells have also been found to possess stem-like characteristics and are resistant to standard-of-care chemotherapy^9^. Targeting stem-like features of RMS would therefore provide novel therapeutic avenues for treating RMS disease relapse and metastasis.

Therapeutic targeting of protein kinases has been demonstrated to be an effective treatment option for a variety of cancers^10^. There exists at least 500 kinases in the human genome, many of which have been linked to the promotion of cancer progression and relapse^10,11^. The roles of kinases in the pathogenesis of cancer and other human diseases have been studied extensively over the past 20 years^12^. However, there currently exists only 48 FDA-approved kinase inhibitors, many of which share the same targets^12^. Of the 48 FDA-approved kinase inhibitors, none have been tested for their therapeutic affects against advanced RMS disease^12^. While previous studies have shown MEK, CDK4/6 and WEE1 as promising kinase targets for inhibiting tumor growth, druggable kinases against RMS self-renewal have been poorly characterized^13,14^. The study by Chen et al (2014) shows that chemical inhibition of glycogen synthase kinase 3 (GSK3) reduces ERMS tumor growth and self-renewal, demonstrating the therapeutic potential for targeting protein kinases that play a role in the regulation of RMS tumor growth and self-renewal^15^.

G-protein coupled receptor kinase 5 (*GRK5)* belongs to a family of serine/threonine kinases^16^ and plays an important role in cardiovascular disease pathogenesis and early heart development^17–19^. GRK5 targets the β-adrenergic receptors, members of the G-protein coupled receptors family (GPCRs), leading to their desensitization and down regulation in cardiomyocytes^20^, and is upregulated during heart failure^21^. GRK5 can also function in a non-GPCR-dependent manner to regulate HDAC5 activity in cardiomyocytes, promoting maladaptive hypertrophy and heart failure^22^. While GRK5 has been extensively studied for its role in heart disease, the role of GRK5 in cancer pathogenesis is poorly characterized. To date, GRK5 has been shown to play a role in the pathogenesis of lung, brain and prostate cancer^4,23,24^. In non-small cell lung cancer (NSCLC) and glioblastoma multiforme (GMB), GRK5 is highly expressed in primary patient specimens and depletion of GRK5 results in reduced cell growth^4,23^. Loss of GRK5 in NSCLC and prostate cancer cell lines also results in cell cycle arrest^23,24^. However, the role of GRK5 in RMS pathogenesis is unknown. GRK5 possesses a unique combination of kinase activity and non-enzymatic protein domains for interacting with substrates, making it an attractive target for drug design in translational applications^20,25,26^.

In this study, we have identified GRK5 as a novel regulator of RMS self-renewal in a high-throughput siRNA library screen against the human kinome (714 kinases). Using the CRISPR/Cas9-based genetic editing strategy, we show that GRK5 loss-of-function reduces RMS self-renewal capacity *in vitro* and *in vivo* through increased programmed cell death. GRK5 regulates cell cycle progression to promote ERMS tumor cell growth in a kinase-independent manner. *NFAT1*, a transcription factor involved in T-cell maturation^27^, is a key player in GRK5-mediated cell cycle progression. Treatment of RMS xenografts with a selective GRK5 inhibitor, CCG-215022, results in a significant reduction of tumor growth, demonstrating the potential of GRK5 as a therapeutic target in RMS.

## Results

### A siRNA library screen of the human kinome identifies GRK5 as a novel regulator of ERMS self-renewal

To identify potential candidate kinases that are essential for self-renewal of ERMS, we performed a siRNA library screen against the human kinome (714 kinases) in two ERMS cell lines (RD and 381T). Each cell line was transfected with a pool of 3 siRNAs against each kinase, along with control (scramble) siRNAs, in 384-well low attachment plates to induce sphere formation. The sphere assay was used as a surrogate *in vitro* assay for assessing the self-renewal capacity of tumor cells^28^. RD and 381T cells were also transfected with the same set of siRNAs in adherent conditions for assessing cell growth. An ATP-based viability assay was performed on siRNA-transfected cells in adherent condition and high-content imaging was performed on the spheres. The normalized ratio of self-renewal capacity to cell growth compared to controls for each kinase target was analyzed (see the volcano plot in Figure 1A). Of the 714 kinases screened, 6 top candidate genes (*FES, LTK, LYN, NME9, PIK3C2A, GRK5*) showed differential effects on self-renewal compared to cell growth and were prioritized for further validation. We subsequently utilized a high-efficiency CRISPR/Cas9 gene targeting strategy^29^ to validate the loss-of-function effect of each candidate kinase gene on self-renewal of 381T ERMS cells (Figure 1B). GRK5 was prioritized for further functional characterization due to consistent loss-of-function effects on self-renewal of RMS cells.

**Figure 1.**
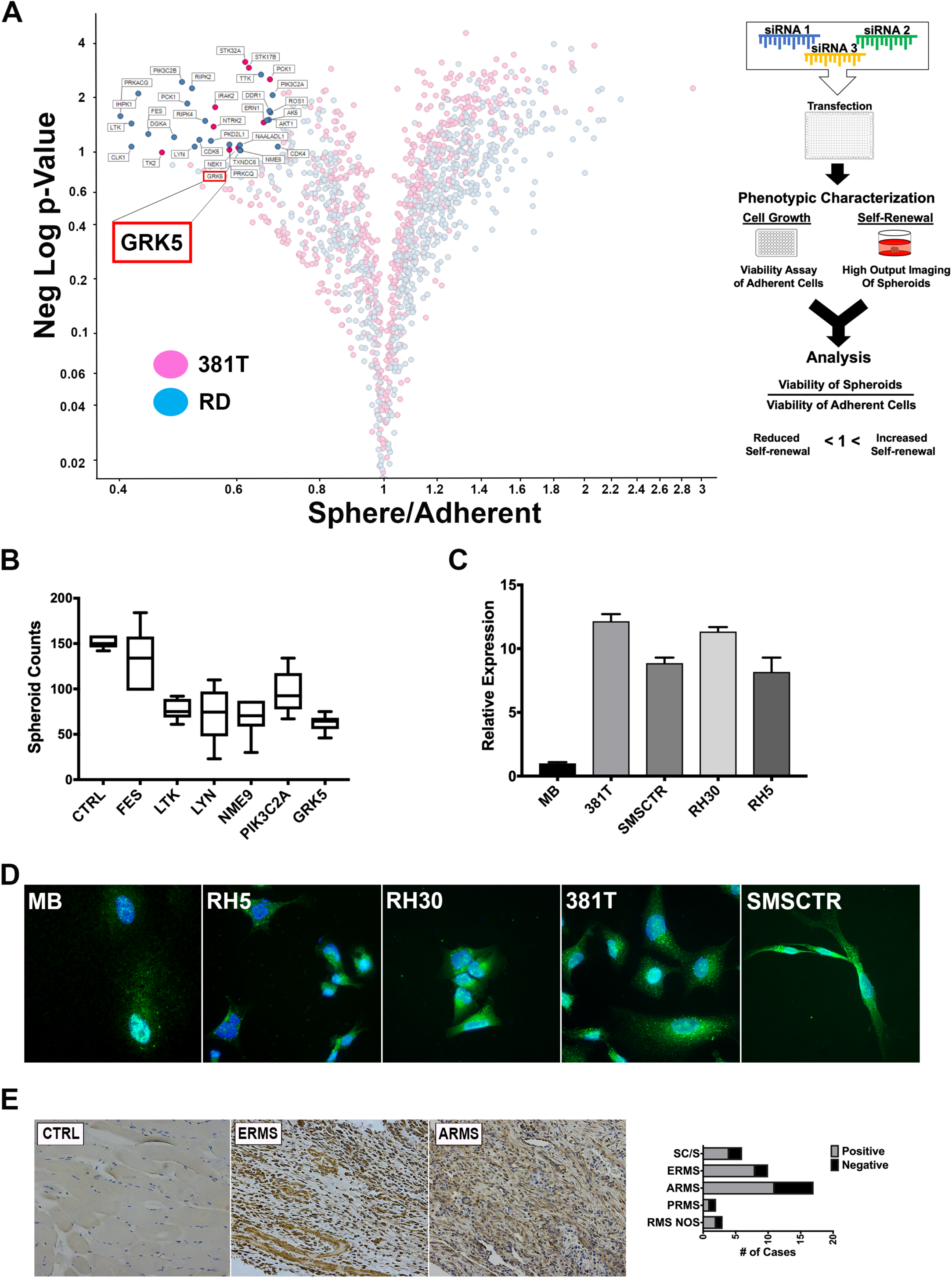
An siRNA library screen of the human kinome identifies GRK5 as a novel regulator of ERMS self-renewal. (A) Volcano plot illustrating candidate kinases identified from an siRNA library screen against the human kinome in ERMS cancer cell lines (381T and RD). Significant hits are indicated as having a p-value (Y-axis) of < 0.05 and a Sphere/Adherent viability ratio (X-axis) of < 1.0. Diagram on the right illustrates workflow and analysis used in the siRNA library screen. GRK5 is highlighted as being a candidate kinase identified from the screen. (B) Spheroid counts to assess self-renewal capacity was performed on CRISPR/Cas9 mediated knockout of top 6 candidate kinases (*FES, LTK, LYN, NME9, PIK3C2A, GRK5)*. Error bars represent standard deviation of 3 technical replicates from an individual experiment that was repeated 3 times. (C) RT-PCR analysis of *GRK5* expression in human myoblasts (MB) compared to a panel of RMS cancer cell lines (381T, SMS-CTR, RH30, RH5). (D) Immunofluorescence images showing GRK5 staining in MB and RMS cancer cell lines (381T, SMSCTR, Rh30, Rh5). (E) Immunohistochemistry of GRK5 in skeletal muscle control (CTRL) and representative primary ERMS and ARMS tumors. Summary of IHC for GRK5 in primary ERMS and ARMS tumors of a tissue microarray is shown on the right.

### GRK5 is differentially expressed in RMS cells compared to normal tissue types and is present in both nuclear and cytoplasmic compartments

*GRK5* mRNA expression levels were analyzed in 4 RMS cell lines (381T and SMSCTR of the ERMS subtype; Rh5 and Rh30 of the ARMS subtype) and compared against a primary myoblast line and an immortalized fibroblast line. In the 4 RMS cell lines, regardless of subtype, the expression level of *GRK5* is at least 2-fold higher compared to normal cell types (Figure 1C).

Immunofluorescence showed both nuclear and cytoplasmic localization of GRK5 in RMS cells (Figure 1D). Immunohistochemistry performed on a tissue microarray (TMA) of primary human RMS tumors showed positive GRK5 expression in the majority of RMS samples including 8/10 ERMS and 10/17 ARMS samples (Figure 1E). In contrast, normal muscle samples from 4 patients showed very weak or negative GRK5 expression. From these findings, *GRK5* appears to be differentially expressed in RMS tumors and likely plays an important role in RMS pathogenesis.

### GRK5 regulates self-renewal of both ERMS and ARMS

To confirm effective disruption of *GRK5* by CRISPR/Cas9, gRNAs were designed to flank the catalytic, nuclear export (NES) and nuclear localization (NLS) functional domains of GRK5 (Figure 2A). Genetic disruption of *GRK5* was then verified via PCR amplification of the genomic deletion event, and depletion of the protein product was confirmed by Western blots (Figure 2A, B). We assessed the loss-of-function effect of GRK5 on tumor cell growth using an ATP-based viability assay on a panel of ARMS (Rh5 and Rh30) and ERMS (381T and SMS-CTR) cell lines. CRISPR/Cas9-mediated disruption of *GRK5* resulted in a significant reduction (p-value < 0.05) in RMS self-renewal capacity via spheroid assay in 3 (381T, SMS-CTR, Rh5) out of the 4 RMS cancer cell lines (Figure 2C). Targeted disruption of *GRK5* in Rh30 showed a trend of reduced self-renewal capacity but was not statistically significant (p-value = 0.086). Spheroids generated from cells harboring targeted disruption of *GRK5* appeared to be both smaller and fewer in number (Figure 2C, D). To determine the cellular mechanism underlying the loss-of-function effects of GRK5 on sphere formation, we performed a quantitative flow cytometry-based Annexin V assay to assess for any change in apoptosis. Loss of GRK5 resulted in a significant increase in early apoptotic events in the spheroids (Figure 2E, F), and approximately 2-fold increase in total apoptotic cells (Figure 2G) compared to the controls. Increased in cell death in GRK5-deficient spheroid cells was further supported by elevated levels of cleaved caspase 3 (CC3) protein (Figure 2H). To assess the effects of GRK5 loss-of-function on the self-renewal capacity of RMS cells *in vivo*, we performed limiting dilution experiments of ERMS (381T) and ARMS (Rh5) xenografts in immunocompromised NOD-scid-IL2Rgammanull (NSG) mice. In both RMS subtypes, targeted disruption of *GRK5* resulted in approximately 4 to 6-fold reduction in self-renewal frequency (Table 1). Taken together, our results indicate that loss of GRK5 in RMS cells results in decreased self-renewal capacity in part through induction of programmed cell death.

**Table 1.** Summary of limiting dilution experiments. 381T and Rh5 RMS cells were injected into NSG mice at limiting dilutions of 2×10^3^, 1×10^4^, 5×10^4^ cells. TPC frequency was calculated as previously described^43^. Confidence interval and statistics was performed on an n = 6 mice per control and GRK5 KO. * = p <0.05; ** = p < 0.01.

| **381T** |  |  |
| --- | --- | --- |
| Cell No. | Control | GRK5 KO |
| 50,000 | 6 of 6 | 6 of 6 |
| 10,000 | 6 of 6 | 5 of 6 |
| 2,000 | 5 of 6 | 3 of 6 |
| **TPC frequency** | 1115 | 4373* |
| 95% CI | 414-3002 | 1927-9922 |

\* p = 0.0353

| **Rh5** |  |  |
| --- | --- | --- |
| Cell No. | Control | GRK5 KO |
| 50,000 | 6 of 6 | 6 of 6 |
| 10,000 | 6 of 6 | 4 of 6 |
| 2,000 | 5 of 6 | 3 of 6 |
| **TPC frequency** | 1115 | 6230** |
| 95% CI | 414-3002 | 2731-14214 |

\*\* p = 0.00739

**Figure 2.**
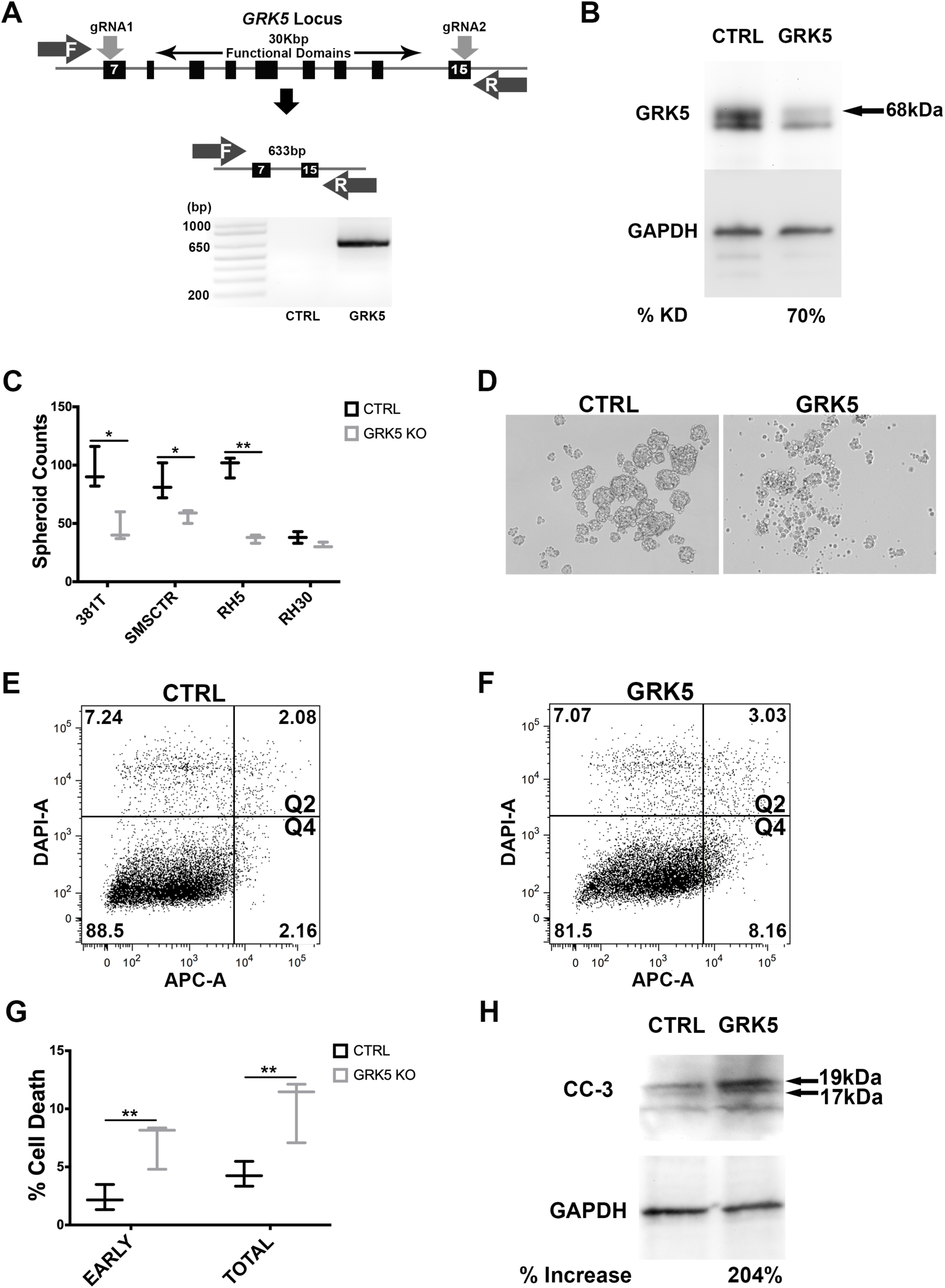
GRK5 regulates self-renewal of both ERMS and ARMS. (A) Schematic demonstrating CRISPR/Cas9 targeting of *GRK5*. Two gRNAs were designed to flank the region encoding functional domains of GRK5. Genetic disruption was confirmed via PCR amplification of the deleted *GRK5* domain followed by Western analysis of depleted protein (B). (C-D) Spheroid counts to assess self-renewal. Results shown are of 3 replicates from one of 3 independent experiments. (E-G) Annexin V Flow cytometry to assess apoptosis of spheroid cells in GRK5 knockout cells (GRK5 KO) compared to Cas9 only controls (CTRL). (H) Western analysis of cleaved caspase 3 (CC3) comparing GRK5 KO spheroids to controls (CTRL). Error bar represents standard deviation and statistical analysis (C, G) was performed on 3 replicates from one of three independent experiments. * = p <0.05; ** = p < 0.01.

### GRK5 regulates ERMS cell growth in a kinase-independent manner and is involved in regulating cell cycle progression

We assessed the loss-of-function effect of GRK5 on tumor cell growth using an ATP-based viability assay on a panel of ARMS (Rh5 and Rh30) and ERMS (381T and SMS-CTR) cell lines. Loss of GRK5 resulted in a significant reduction in cell viability in all 4 RMS cell lines (p-value < 0.05) (Figure 3A). To assess the specificity of GRK5 loss-of-function effect on RMS cell growth, we overexpressed a Cas9-resistant form of GRK5 in the presence of CRISPR/Cas9-mediated *GRK5* gene disruption in SMS-CTR cells. Compared to GFP overexpression control, Cas9-resistant GRK5 rescued the growth phenotype of SMS-CTR cells following targeted disruption of *GRK5* (Figure 3B). Even though the GRK family proteins are known for their kinase-dependent roles, some studies have also implicated kinase-independent function of GRK5^20,30^. To determine whether GRK5 regulates RMS cell growth in a kinase dependent or independent manner, we generated a kinase dead (K215R) (KD) GRK5 mutant^25^ that is resistant to targeted disruption by CRISPR/Cas9. Overexpression of KD GRK5 protein also restored cell growth in SMS-CTR cells with targeted disruption of *GRK5* (Figure 3B), indicating that GRK5 regulates ERMS cell growth in a kinase-independent manner. We next determined the cellular event that was responsible for GRK5 loss-of-function effect on RMS cell growth. While 381T and SMS-CTR cells with targeted disruption of *GRK5* showed no significant change in cellular differentiation or cell death (Figure 3C, D), they showed altered cell cycle progression in a flow cytometry-based cell cycle analysis following EdU pulse (Figure 3E, F). There was an arrest in the G1/S phase in 381T cells, and in the G2/M phase in SMS-CTR cells. Overall, our results indicate that GRK5 functions in a kinase-independent manner to alter cell cycle progression in ERMS cells.

**Figure 3.**
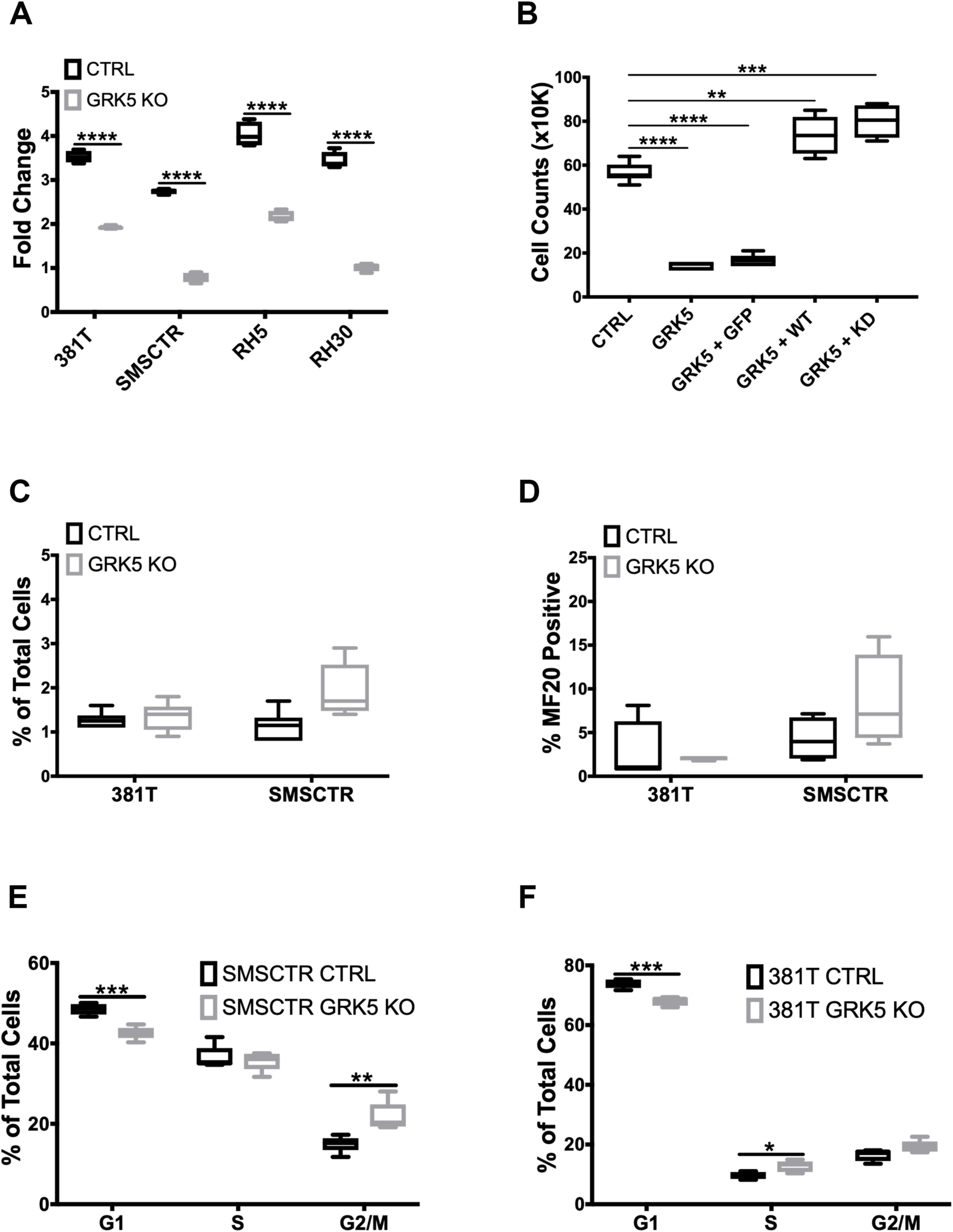
GRK5 regulates ERMS cell growth in a kinase-independent manner and is involved in regulating cell cycle progression. (A) Cell Titer Glo viability assessment of *GRK5* knockout (GRK5 KO) compared to controls (CTRL) in a panel of RMS cancer cell lines (381T, SMS-CTR, RH5, RH30). Data shown are 4 replicates from one of 3 independent experiments, **** = p < 0.0001. (B) Summary of cell count data from rescue experiment to demonstrate specificity of the *GRK5* KO growth phenotype. SMSCTR cells received lentivirus containing either Cas9 only (CTRL), Cas9 with *GRK5* gRNAs (GRK5), Cas9 with *GRK5* gRNAs and GFP overexpression (GRK5 + GFP), Cas9 with *GRK5* gRNAs and Cas9 resistant, wildtype GRK5 (GRK5 + WT), Cas9 w/ *GRK5* gRNAs and Cas9 resistant, kinase dead GRK5 (GRK5 + KD). Data shown are 6 replicates from one of three independent experiments. ** = p < 0.01, *** = p < 0.001, (C) Annexin V Flow cytometry to assess apoptosis. (D) Quantitation of immunofluorescence (IF) against MF20 in RMS cells with GRK5 knockout (GRK5 KO) and Cas9 only controls (CTRL). (E-F) EdU flow cytometry-based cell cycle analysis of SMSCTR and 381T cells with Cas9 only as controls (CTRL) or GRK5 knockout (GRK5 KO). Data shown are from 6 independent experiments. * = p < 0.05, ** = p < 0.01, *** = p < 0.001, **** = p < 0.0001.

### NFAT1 is a key player in GRK5 mediated cell cycle progression

GRK5 has been previously shown to facilitate the transcriptional activity of NFAT as part of a DNA binding complex during cardiac hypertrophy in a kinase independent manner^20^. NFAT1 has been shown to function either as a positive or negative regulator of cell cycle progression^31,32^. To investigate the role of NFAT as a potential downstream component of the GRK5 pathway in regulating ERMS cell growth, we first showed that targeted disruption of *GRK5* led to decreased levels of NFAT1 expression in 381T cells (Figure 4A). Overexpression of GRK5 restored expression levels of *NFAT1* (Figure 4B). To determine the loss-of-function effects of NFAT1 on RMS cells, we showed that targeted disruption of *NFAT1* by CRISPR/Cas9 in ERMS cells significantly reduced cell growth (Figure 4C). Cell cycle analysis of *NFAT1*-targeted SMS-CTR cells showed significant alteration of cell cycle progression (Figure 4D). There was an arrest in both G1/S and G2/M in *NFAT1-*disrupted SMS-CTR cells. To determine if GRK5 interacts with NFAT1 in ERMS, we showed by immunofluorescence that GRK5 and NFAT1 co-localized in 381T cells (Figure 4E). By proximity ligation assay, we showed direct interaction of GRK5 and NFAT in the nucleus and the cytoplasm of 381T cells, and that loss of GRK5 abrogated this interaction (Figure 4F). Taken together, our data indicate that NFAT1 is a key mediator of GRK5 function in regulating RMS cell growth.

**Figure 4.**
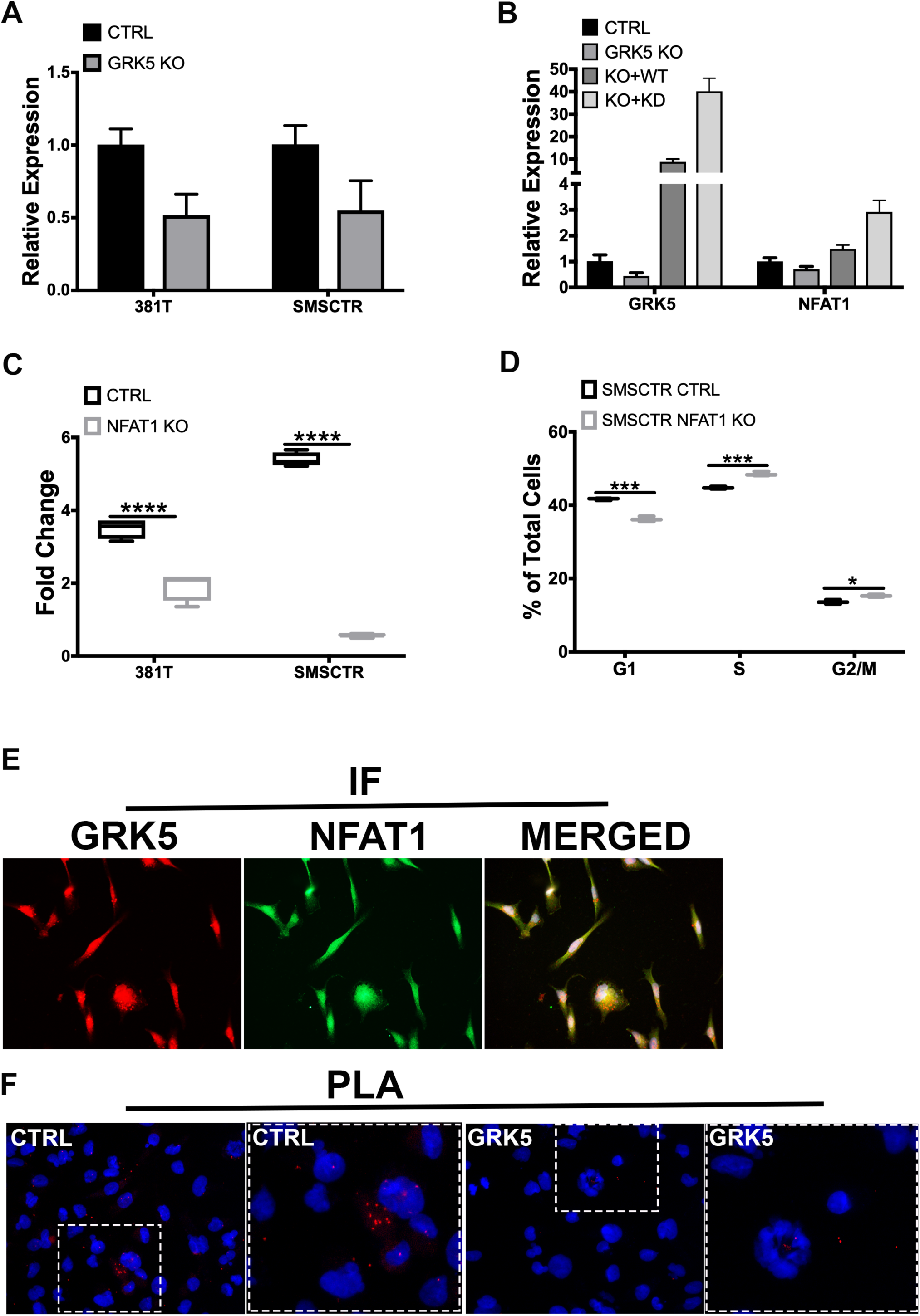
NFAT1 is a key player in GRK5-mediated cell cycle progression. (A) RT-PCR analysis of *NFAT1* expression in ERMS cells (381T/SMS-CTR) with GRK5 (CTRL) and GRK5 knockout (GRK5 KO). (B) RT-PCR analysis of *GRK5* and *NFAT1* expression in 381T cells following GRK5 rescue with wild-type and domain mutants. Cas9 only (CTRL), Cas9 with *GRK5* gRNAs (GRK5), Cas9 with *GRK5* gRNAs and Cas9 resistant, wildtype GRK5 (GRK5 + WT), Cas9 with *GRK5* gRNAs and Cas9 resistant, kinase dead GRK5 (GRK5 + KD). (C) Cell Titer Glo viability assessment of *NFAT1* knockout (NFAT1 KO) compared to controls (CTRL) in ERMS cancer cell lines (381T, SMS-CTR). Data shown are 4 replicates from one of 3 independent experiments, **** = p < 0.0001. (D) EdU flow cytometry-based cell cycle analysis of SMS-CTR cells with Cas9 only as controls (CTRL) or *NFAT1* knockout (NFAT1 KO). Data shown are of 3 replicates from one of 3 independent experiments. * = p < 0.05, *** = p < 0.001. (E) Immunofluorescence images showing GRK5 (red) and NFAT1 (green) staining in 381T cells, with overlay of both channels (yellow). (F) Proximity Ligation Assay (PLA) to assess GRK5-NFAT1 protein interaction in GRK5 wildtype (CTRL) and GRK5 deficient (GRK5) 381T cells. Red dots represent points of GRK5-NFAT1 close proximity.

### Treatment of RMS tumor with CCG-215022, a GRK5 inhibitor, reduces tumor growth *in vivo*

To assess the potential of GRK5 as a therapeutic target against RMS, 381T or Rh5 RMS xenografts established in NSG mice were treated with a GRK5-selective inhibitor, CCG-215022 (Figure 5A). Mice treated with CCG-215022 showed a significant reduction in tumor growth in both 381T and Rh5 tumors compared to the mice treated with vehicle control (Figure 5B-D). Immunohistochemistry analysis of CCG-215022 treated tumors shows a lower Ki-67 proliferation index compared to the vehicle control-treated tumors. These results demonstrate the therapeutic potential of targeting GRK5 as an alternative treatment option to inhibit RMS tumor growth.

**Figure 5.**
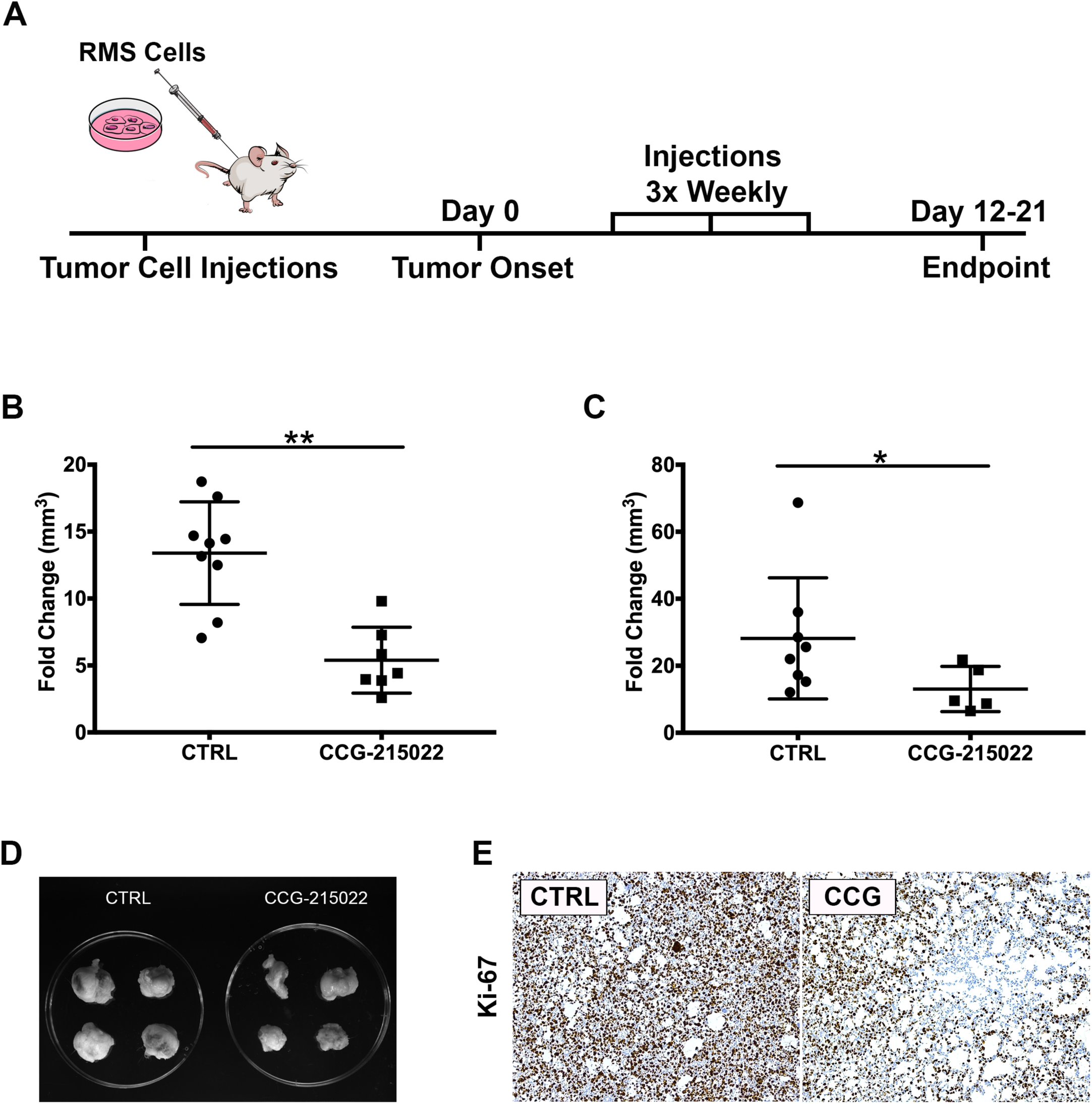
Treatment of RMS tumor with CCG-215022, a GRK5 inhibitor, reduces tumor growth *in vivo.* (A) Schematic of the RMS xenograft in NSG mice experiment. (B-C) Summary of 381T (B) and Rh5 (C) tumor volume fold change following CCG-215022 or DMSO control (CTRL) treatment. Each data point represents one mouse with error bars representing standard deviation. * = p < 0.05, ** = p < 0.01 (Mann-Whitney Statistical Test). (C-D) Representative images of Rh5 tumors harvested from DMSO vehicle control (CTRL) treated and CCG-215022 treated mice at the end of treatment period (D). (E) Representative images of Immunohistochemistry for Ki67 on DMSO control (CTRL) and CCG-215022 (CCG)-treated tumors.

## Discussion

While *GRK5* has been extensively studied for its role in the pathogenesis of cardiovascular disease, only in recent years has its role in cancer biology been brought to light^17–22^. GRK5 has been implicated as having a role in regulating lung, brain and prostate cancer cell growth^4,23,24^. However, the role of GRK5 in RMS has yet to be investigated. In this study, we show that GRK5 is a novel regulator of RMS cell growth through its interaction with NFAT1 to regulate cell cycle progression in a kinase-independent manner. Loss of GRK5 reduces the self-renewal capacity of RMS cells through increased programmed cell death, implicating it as a novel regulator of stem-like features in RMS. We demonstrate the potential for GRK5 as a therapeutic target through treatment of RMS xenografts with a selective GRK5 inhibitor, CCG-215022. RMS tumors treated with CCG-215022 show significant reduction in tumor growth and self-renewal capacity.

The studies characterizing the role of GRK5 in cancer pathogenesis to date have primarily investigated the functional requirement of its kinase activity^33,34^. TP53 phosphorylation by GRK5 leads to its degradation, resulting in inhibition of the TP53-dependent apoptotic response to genotoxicity in osteosarcoma cells^33^. A study using HeLa cells additionally shows a defect in proper cell cycle progression following *GRK5* gene knockdown^34^. However, we show through rescue experiments with a GRK5 kinase-dead mutant that GRK5 regulates RMS cell growth in a kinase-independent manner. We also show that targeted disruption of *GRK5* in two RMS cell lines with TP53 mutations, 381T with *TP53*(*R248W*) and Rh30 with *TP53*(*R237H)*^35,36^, reduces RMS cell growth by inducing cell cycle arrest. Our findings indicate that GRK5 regulates RMS cell growth in a kinase- and TP53-independent manner.

Dysregulated activity of the NFAT family of transcription factors has been identified in cancer^37^. While NFAT proteins (NFAT1-5) share similar DNA binding targets, each member possesses both redundant and opposing functions, and their activity often requires cooperation with additional transcriptional partners^31,32,37^. NFAT1 has been shown to function both as a tumor suppressor through the transcriptional activation of the CDK4 promoter or as an oncogene by silencing p15 expression^20^. In a model of pathological cardiac hypertrophy, GRK5 promotes the transcriptional activity of NFAT in a kinase-independent manner as part of a DNA binding complex to regulate expression of hypertrophic genes^20^. In this study, both wild-type and kinase-dead GRK5 are able to restore *NFAT1* expression following targeted disruption of *GRK5* in ERMS cell lines. We also show that GRK5 directly interacts with NFAT1, and this interaction is abrogated with loss of GRK5. Our findings indicate a novel role for GRK5 acting in a kinase-independent manner to facilitate the autoregulation of *NFAT1* expression. CRISPR/Cas9 mediated disruption of NFAT1 phenocopies the loss-of-function effects of GRK5 on cell cycle progression. Together, our findings demonstrate that the interaction between GRK5 and NFAT1 is essential for ERMS tumor cell growth.

TPCs undergo self-renewal to recapitulate the complex heterogeneity of a given malignant tumor and are thought to be the major drivers of cancer relapse and metastasis in selected cancer types^7,38^. Disease relapse or metastasis of RMS carries a poor survival prognosis^3^. Identifying potential targets that regulate TPC survival could potentially provide a solution for treating RMS disease relapse and metastasis. Our study shows that GRK5 loss-of-function significantly reduces the self-renewal capacity of RMS cells *in vitro* and *in vivo*. The stem-like RMS spheres harboring *GRK5* knockout show increased cell death. Our findings indicate that GRK5 is a promising therapeutic target against RMS stem-like features. Further investigation is required to assess whether loss of GRK5 leads to reduced heterogeneity of RMS tumors and thereby reduces the potential for resistance against standard-of-care therapies.

Inhibitors selective to GRK5 are currently limited. Amlexanox, an FDA-approved anti-inflammatory drug, has been shown to inhibit GRK5 activity^39^. However, amlexanox is a non-specific inhibitor with cross reactivity with other proteins and pathways such as IKBKE in the Hippo pathway^40^. CCG-215022, an investigational compound developed by John Tesmer’s group at the University of Michigan, shows high selectivity against GRK5^41^. In our study, treatment of both ERMS and ARMS xenograft tumors treated with CCG-215022 significantly reduces tumor growth. CCG-215022-bound GRK5 shows disorder in the residues pertaining to the N-terminus, C-terminus and active site tether regions^41^. We show that a kinase deficient GRK5 mutant rescues the GRK5 loss-of-function growth phenotype. Based on this finding, it is possible that the functional domain of GRK5 that regulates RMS cell growth likely resides in one of these disordered regions. While additional testing to assess the toxicity profile of CCG-215022 in pre-clinical models is necessary, we have shown that inhibition of GRK5 is a promising therapeutic option for RMS patients.

With treatment options against RMS remaining relatively unchanged over last 3 decades, there remains a need for more effective therapeutic targets. From a comprehensive siRNA library screen against the human kinome, we have identified GRK5 as a novel regulator of both RMS self-renewal and cell growth. Our functional characterization of GRK5 *in vitro* and *in vivo* demonstrates that GRK5 regulates ERMS cell growth in a kinase-independent manner and is essential for RMS self-renewal capacity. A GRK5 inhibitor, CCG-215022, recapitulates the loss-of-function effects of GRK5. Thus, our findings demonstrate the promise of GRK5 as a therapeutic target against RMS disease progression and relapse.

## Methods

### siRNA kinome library screen

To identify potential candidate kinases that are important for the self-renewal of ERMS, we utilized the Quellos high-throughput screening core facility in the Institute of Stem Cell and Regenerative Medicine at the University of Washington to perform an siRNA library screen against the human kinome (714 kinases) in two different cell lines derived from ERMS (RD and 381T). Each cell line was transfected with a pool of 3 siRNAs against each kinase, along with control (scramble) siRNAs, in 384-well low attachment plates to induce sphere formation. The sphere assay was used as a surrogate *in vitro* assay for assessing the self-renewal capacity of tumor cell. The ATP-based Cell Titer Glo assay (Promega) was performed on the siRNA-treated adherent cells, and high-content imaging was performed at the Quellos core facility at day 5 post-siRNA transfection.

### CRISPR/Cas9 gene targeting in human RMS cells

Single gene knockout was accomplished using lentiviral transduction of RMS cells with Cas9 expressing and gene-specific double gRNA constructs. Lentiviral transduced cells were placed under antibiotic selection and plated for assays 7 days later. Cloning of Cas9 and gRNA expression constructs was performed as described previously^36^. Overexpression constructs used in GRK5 functional experiments were amplified from cDNA generated from ERMS cancer cell RNA. Silent mutations to PAM sites were introduced to GRK5 overexpression constructs to generate Cas9 resistant GRK5 protein. RMS cells were then transduced with 3 separate viruses; Cas9 virus, dgRNA virus and GRK5 WT/KD overexpression virus. Domain mutations made to GRK5 to generate wildtype (WT) and kinase dead (KD) variants was done using Gibson cloning based off of previous studies^25^.

The following gRNAs were used for targeting genes in human RMS cell lines:

*GRK5:* gRNA1- GGACCTGGTCTCCCAGACGG

gRNA2- GGAGCAGCCCTTTCTTGGG

*NFAT1: gRNA1- GACGGAGTGATCTCGATCCG*

*gRNA2- GATCCCACAAGGCGAGTCCG*

*FES:* gRNA1-GGCCGAGCTTCGTCTACTGG

gRNA2- GAGCCTGCTCATCCGGGAA

*LTK:* gRNA1- GCTGGCTCCAAGATACTAGG

gRNA2- GACCAGCGTGGTGGTGACCG

*LYN:* gRNA1- GTAGCCTTGTACCCCTATGA

gRNA2- GGAATGGCATACATCGAG

*NME9:* gRNA1- GACCTCGATCCTCATCTTC

gRNA2- GATGTCCTCGAAAAGTACAG

*PIK3C2A:* gRNA1- GCACAGGTTTATAACAAGC

gRNA2- GGGGCGCTTGCTAATATTTT

### Cell-based Assays

RMS cell growth was assessed via cell counts or an ATP-based luminescent cell viability assay,CellTiter-Glo (Promega, Madison, WI). Myogenic differentiation was performed following serum starvation of RMS cells in 2% horse serum/DMEM for 72 hours prior to fixation in 2% paraformaldehyde. Immunofluorescence against MF20 (myosin heavy chain) was then performed. Self-renewal was assessed by visual counts of rhabdospheres, induced in growth factors (EGF, bFGF, PDGF-A, PDGF-B) enriched neurobasal medium as previous described^42^. Apoptosis was assessed using a flow-cytometry based assay using the Annexin V, Alexa Fluor 647 conjugate (Life Technologies). Cell cycle analysis was done with flow cytometry on cells pulsed with EdU for 2 hours using the Click-iT EdU Alexa Fluor 647 Flow Cytometry Assay kit (Life Technologies, Carlsbad, CA).

### Human Xenografts and Drug Treatment

All mouse experiments were approved by the University of Washington Subcommittee on Research Animal Care under IACUC protocol #4330-01. 6-7 immunocompromised NOD-SCID IL2rg-/-(NSG) mice were xenografted via subcutaneous injections into the flanks with approximately 1-2×10^6^ RMS cells (RH5 or 381T) suspended in Matrigel. At tumor onset, CCG-215022 (10mg/kg) or vehicle (DMSO) were given intraperitoneal every 3 days for 21 days or until tumors reached end point. Tumor measurements were made with calipers every 3-4 days at tumor onset until tumors reached end point or at the end of drug treatment, whichever came first. For limiting dilution experiments, 6 NSG mice were given RMS cell injections (Rh5 or 381T) of either control or *GRK5* knock-out cells at either 2×10^3^, 1×10^4^, 5×10^4^ dilutions in the same manner as previously described. Analysis of limiting dilution data was performed as previously described^43^. All mice were humanly euthanized for tumor tissue harvesting at the end of the experiment.

### Immunohistochemistry and Immunofluorescence

The RMS tissue microarray was obtained from Seattle Children’s Hospital. Immunohistochemistry was performed at the Histology and Imaging core facility at the University of Washington. Immunofluorescence was performed as previously described^29^. Duolink Proximity Ligation Assay (PLA) by Sigma Aldrich was used to visualize GRK5-NFAT1 activity. The following antibodies were used: rabbit polyclonal anti human mouse monoclonal anti human Ki-67, 1:100,(MIB1), Dako), Immunohistochemistry: GRK5 1:100 (N145/20) Abcam, Cambridge, MA), Immunofluorescence: GRK5 1:150, (N145/20), Abcam, Cambridge, MA), NFAT1 1:100, (D43B1), Cell Signaling Technology, Danvers, MA)

### Western Blots

Cell lysates from RMS cells were counted and lysed in RIPA buffer (ThermoFisher #89900) with protease inhibitors plus 2x sample buffer (100mM Tris pH6.8, 4%SDS, 20% glycerol). Equal amounts of protein lysates were electrophoresed on a 4-15% gradient SDS-polyacrylamide gel (BioRad, Hercules, CA) and fast transferred to Immun-Blot PVDF membranes (BioRad, Hercules, CA) using the Turbo-Blot Transfer system (BioRad, Hercules, CA) We used the following antibodies: GRK5 (N145/20) 1:100 Abcam, Cambridge, MA), GAPDH (14C10) 1:2500 (Cell Signaling Technology, Danvers, MA), Cleaved Caspase 3 (Asp175) 1:200 (Cell Signaling Technology, Danvers, MA). Goat anti-mouse or anti-rabbit HRP conjugated IgG secondary antibodies from Santa Cruz Biotechnology. Membranes were blocked in 5% milk in Tris Buffered Saline plus Tween (20mM Tris, 136mM NaCl, 1% Tween 20, pH 7.4 TBST). Quantitative analysis of Western blot images was performed on ImageJ.

### Human Expression Data Analysis

RNA was collected from human cell lines lysates (myblasts, NHDF, 381T, SMSCTR, RH30, RH5) using Qiagen RNeasy Plus Mini Kit. cDNA was then generated using High Capacity cDNA Reverse Transcription Kit from Applied Biosystems. RT-PCR reactions were then run with iTaq Universal SYBR Green mix on a CFX Connect Real Time System (BioRad, Hercules, CA). RT-PCR primers used are listed below:

*GRK5* FWD – GTCTGTCCACGAGTACCTGA

REV – CAGGCATACATTTTACCCGT

*NFAT1* FWD – ACGAGCTTGACTTCTCCACC

REV – TGCATTCGGCTCTTCTTCGT

### Statistics

Mann-Whitney statistical test was run on drug treat RMS tumor mouse experiments to assess statistical significance in differences between experimental and control. Two tailed Student’s t-test was applied when appropriate.

## Acknowledgements

We would like to thank the Molecular Medicine and Mechanism of Disease (M3D) PhD program at the University of Washington School of Medicine for their support. Additional thanks to all the undergraduates and Chen Lab members who provided technical assistance to the study (Phuong Van, Shelly Lin, Henna Di, Dr. Michael Phelps, Maria Pavlova and Dr. Julia Sidorova). EYC is supported by NIH R01 CA196882.

## References

1. Shern, J. F. et al. Comprehensive genomic analysis of rhabdomyosarcoma reveals a landscape of alterations affecting a common genetic axis in fusion-positive and fusion-negative tumors. Cancer Discov 4, 216–231 (2014).

2. Barr, F. G. et al. Rearrangement of the PAX3 paired box gene in the paediatric solid tumour alveolar rhabdomyosarcoma. Nat Genet 3, 113–117 (1993).

3. Pappo, A. S. et al. Survival After Relapse in Children and Adolescents With Rhabdomyosarcoma: A Report From the Intergroup Rhabdomyosarcoma Study Group. JCO 17, 3487–3493 (1999).

4. Kaur, G. et al. G-protein coupled receptor kinase (GRK)-5 regulates proliferation of glioblastoma-derived stem cells. Journal of Clinical Neuroscience 20, 1014–1018 (2013).

5. Lawson, D. A. et al. Single-cell analysis reveals a stem-cell program in human metastatic breast cancer cells. Nature 526, 131–135 (2015).

6. Jahchan, N. S. et al. Identification and targeting of long-term tumor-propagating cells in small cell lung cancer. Cell Rep 16, 644–656 (2016).

7. Kreso, A. & Dick, J. E. Evolution of the Cancer Stem Cell Model. Cell Stem Cell 14, 275–291 (2014).

8. Ignatius, M. S. et al. In Vivo Imaging of Tumor-Propagating Cells, Regional Tumor Heterogeneity, and Dynamic Cell Movements in Embryonal Rhabdomyosarcoma. Cancer Cell 21, 680–693 (2012).

9. Walter, D. et al. CD133 Positive Embryonal Rhabdomyosarcoma Stem-Like Cell Population Is Enriched in Rhabdospheres. PLOS ONE 6, e19506 (2011).

10. Gross, S., Rahal, R., Stransky, N., Lengauer, C. & Hoeflich, K. P. Targeting cancer with kinase inhibitors. J Clin Invest 125, 1780–1789 (2015).

11. Hanahan, D. & Weinberg, R. A. Hallmarks of Cancer: The Next Generation. Cell 144, 646–674 (2011).

12. Roskoski, R. Properties of FDA-approved small molecule protein kinase inhibitors. Pharmacological Research 144, 19–50 (2019).

13. Yohe, M. E. et al. MEK inhibition induces MYOG and remodels super-enhancers in RAS-driven rhabdomyosarcoma. Science Translational Medicine 10, eaan4470 (2018).

14. Stewart, E. et al. Identification of Therapeutic Targets in Rhabdomyosarcoma through Integrated Genomic, Epigenomic, and Proteomic Analyses. Cancer Cell 0, (2018).

15. Chen, E. Y. et al. Glycogen synthase kinase 3 inhibitors induce the canonical WNT/β-catenin pathway to suppress growth and self-renewal in embryonal rhabdomyosarcoma. Proc Natl Acad Sci U S A 111, 5349–5354 (2014).

16. Willets, J. M., Challiss, R. A. J. & Nahorski, S. R. Non-visual GRKs: are we seeing the whole picture? Trends in Pharmacological Sciences 24, 626–633 (2003).

17. Islam, K. N., Bae, J.-W., Gao, E. & Koch, W. J. Regulation of Nuclear Factor κB (NF-κB) in the Nucleus of Cardiomyocytes by G Protein-coupled Receptor Kinase 5 (GRK5). J. Biol. Chem. 288, 35683–35689 (2013).

18. Philipp, M., Berger, I. M., Just, S. & Caron, M. G. Overlapping and Opposing Functions of G Protein-coupled Receptor Kinase 2 (GRK2) and GRK5 during Heart Development. J Biol Chem 289, 26119–26130 (2014).

19. Zhang, Y. et al. Nuclear Effects of GRK5 on HDAC5-regulated Gene Transcription in Heart Failure. Circ Heart Fail 4, 659–668 (2011).

20. Hullmann, J. E. et al. GRK5-Mediated Exacerbation of Pathological Cardiac Hypertrophy Involves Facilitation of Nuclear NFAT Activity. Circ Res 115, 976–985 (2014).

21. Lessel, D. et al. The analysis of heterotaxy patients reveals new loss-of-function variants of GRK5. Scientific Reports 6, 33231 (2016).

22. Martini, J. S. et al. Uncovering G protein-coupled receptor kinase-5 as a histone deacetylase kinase in the nucleus of cardiomyocytes. Proc Natl Acad Sci U S A 105, 12457–12462 (2008).

23. Jiang, L.-P. et al. GRK5 functions as an oncogenic factor in non-small-cell lung cancer. Cell Death Dis 9, (2018).

24. Kim, J. I., Chakraborty, P., Wang, Z. & Daaka, Y. G-Protein Coupled Receptor Kinase 5 Regulates Prostate Tumor Growth. The Journal of Urology 187, 322–329 (2012).

25. Johnson, L. R., Robinson, J. D., Lester, K. N. & Pitcher, J. A. Distinct Structural Features of G Protein-Coupled Receptor Kinase 5 (GRK5) Regulate Its Nuclear Localization and DNA-Binding Ability. PLOS ONE 8, e62508 (2013).

26. Sorriento, D. et al. The Amino-Terminal Domain of GRK5 Inhibits Cardiac Hypertrophy through the Regulation of Calcium-Calmodulin Dependent Transcription Factors. Int J Mol Sci 19, (2018).

27. Macian, F. NFAT proteins: key regulators of T-cell development and function. Nature Reviews Immunology 5, nri1632 (2005).

28. Pastrana, E., Silva-Vargas, V. & Doetsch, F. Eyes Wide Open: A Critical Review of Sphere-Formation as an Assay For Stem Cells. Cell Stem Cell 8, 486–498 (2011).

29. Phelps, M. P., Bailey, J. N., Vleeshouwer-Neumann, T. & Chen, E. Y. CRISPR screen identifies the NCOR/HDAC3 complex as a major suppressor of differentiation in rhabdomyosarcoma. PNAS 113, 15090–15095 (2016).

30. Watari, K., Nakaya, M. & Kurose, H. Multiple functions of G protein-coupled receptor kinases. J Mol Signal 9, 1 (2014).

31. Baumgart, S. et al. Restricted Heterochromatin Formation Links NFATc2 Repressor Activity With Growth Promotion in Pancreatic Cancer. Gastroenterology 142, 388–98.e1–7 (2012).

32. Baumgart, S. et al. Inflammation induced NFATc1-STAT3 Transcription Complex Promotes Pancreatic Cancer initiation by KrasG12D. Cancer Discov 4, 688–701 (2014).

33. Chen, X. et al. G-protein-coupled Receptor Kinase 5 Phosphorylates p53 and Inhibits DNA Damage-induced Apoptosis. J Biol Chem 285, 12823–12830 (2010).

34. Michal, A. M. et al. G Protein-coupled Receptor Kinase 5 Is Localized to Centrosomes and Regulates Cell Cycle Progression. J Biol Chem 287, 6928–6940 (2012).

35. Ognjanovic, S. et al. Low Prevalence of TP53 Mutations and MDM2 Amplifications in Pediatric Rhabdomyosarcoma. Sarcoma https://www.hindawi.com/journals/sarcoma/2012/492086/ (2012) doi:10.1155/2012/492086.

36. Hinson, A. R. P. et al. Human Rhabdomyosarcoma Cell Lines for Rhabdomyosarcoma Research: Utility and Pitfalls. Front Oncol 3, (2013).

37. Mognol, G. P., Carneiro, F. R. G., Robbs, B. K., Faget, D. V. & Viola, J. P. B. Cell cycle and apoptosis regulation by NFAT transcription factors: new roles for an old player. Cell Death Dis 7, e2199 (2016).

38. Saygin, C., Matei, D., Majeti, R., Reizes, O. & Lathia, J. D. Targeting Cancer Stemness in the Clinic: From Hype to Hope. Cell Stem Cell 24, 25–40 (2019).

39. Homan, K. T., Wu, E., Cannavo, A., Koch, W. J. & Tesmer, J. J. G. Identification and Characterization of Amlexanox as a G Protein-Coupled Receptor Kinase 5 Inhibitor. Molecules 19, 16937–16949 (2014).

40. Liu, Y. et al. Amlexanox, a selective inhibitor of IKBKE, generates anti-tumoral effects by disrupting the Hippo pathway in human glioblastoma cell lines. Cell Death & Disease 8, e3022 (2017).

41. Homan, K. T. et al. Crystal Structure of G Protein-Coupled Receptor Kinase 5 in Complex with a Rationally Designed Inhibitor. J. Biol. Chem. jbc.M115.647370 (2015) doi:10.1074/jbc.M115.647370.

42. Vleeshouwer-Neumann, T. et al. Histone Deacetylase Inhibitors Antagonize Distinct Pathways to Suppress Tumorigenesis of Embryonal Rhabdomyosarcoma. PLoS One 10, (2015).

43. Hu, Y. & Smyth, G. K. ELDA: Extreme limiting dilution analysis for comparing depleted and enriched populations in stem cell and other assays. Journal of Immunological Methods 347, 70–78 (2009).

